# Structural neuroplasticity after sleep loss modifies behavior and requires neurexin and neuroligin

**DOI:** 10.1101/2023.06.20.545788

**Authors:** Mara H. Cowen, David M. Raizen, Michael P. Hart

## Abstract

Sleep is conserved across species, and alterations in sleep are associated with a multitude of neurologic conditions. Structural neuroplasticity, or changes in the size, strength, number, and targets of neuronal synaptic connections, can be modified by sleep and sleep disruption. However, the causal relationships between sleep loss and neuroplasticity, behavioral impacts, and genetic perturbations are poorly understood. The *C. elegans* GABAergic DVB neuron undergoes structural plasticity in adult males that rewires synaptic connections and alters behavior; the rewiring is dependent on the conserved autism-associated genes *NRXN1*/*nrx-1* and *NLGN3/nlg-1.* Stress exposure during sexual maturation, which overlaps with a developmental sleep period, alters DVB structural plasticity in early adulthood. Here, we use four distinct methods to ask whether sleep deprivation results in altered structural and behavioral plasticity of the DVB neuron. Following sleep deprivation using the genetic mutants (*aptf-1* and *npr-1*), vibration stimulus, or chemo-genetic silencing of a sleep-producing neuron, we find that DVB neurite outgrowth at day 1 of adulthood. Sleep deprivation also leads to an increase in the time to spicule protraction, the behavioral output of DVB function. We find that *nrx-1* and *nlg-1* both function to regulate sleep loss induced DVB structural and functional changes. Lastly, differences in DVB neurite outgrowth in sleep deprived animals at day 1 are transient, with DVB morphology being indistinguishable from controls by day 3. These results show that sleep loss modifies DVB structural plasticity and its behavioral consequences. Further, we show *nrx-1* and *nlg-1* mediate the effect of sleep deprivation on behavior at the level of DVB structural plasticity. These insights into sleep, neuroplasticity, behavior, and conserved autism genes have important implications for the role of sleep in neurologic disease.

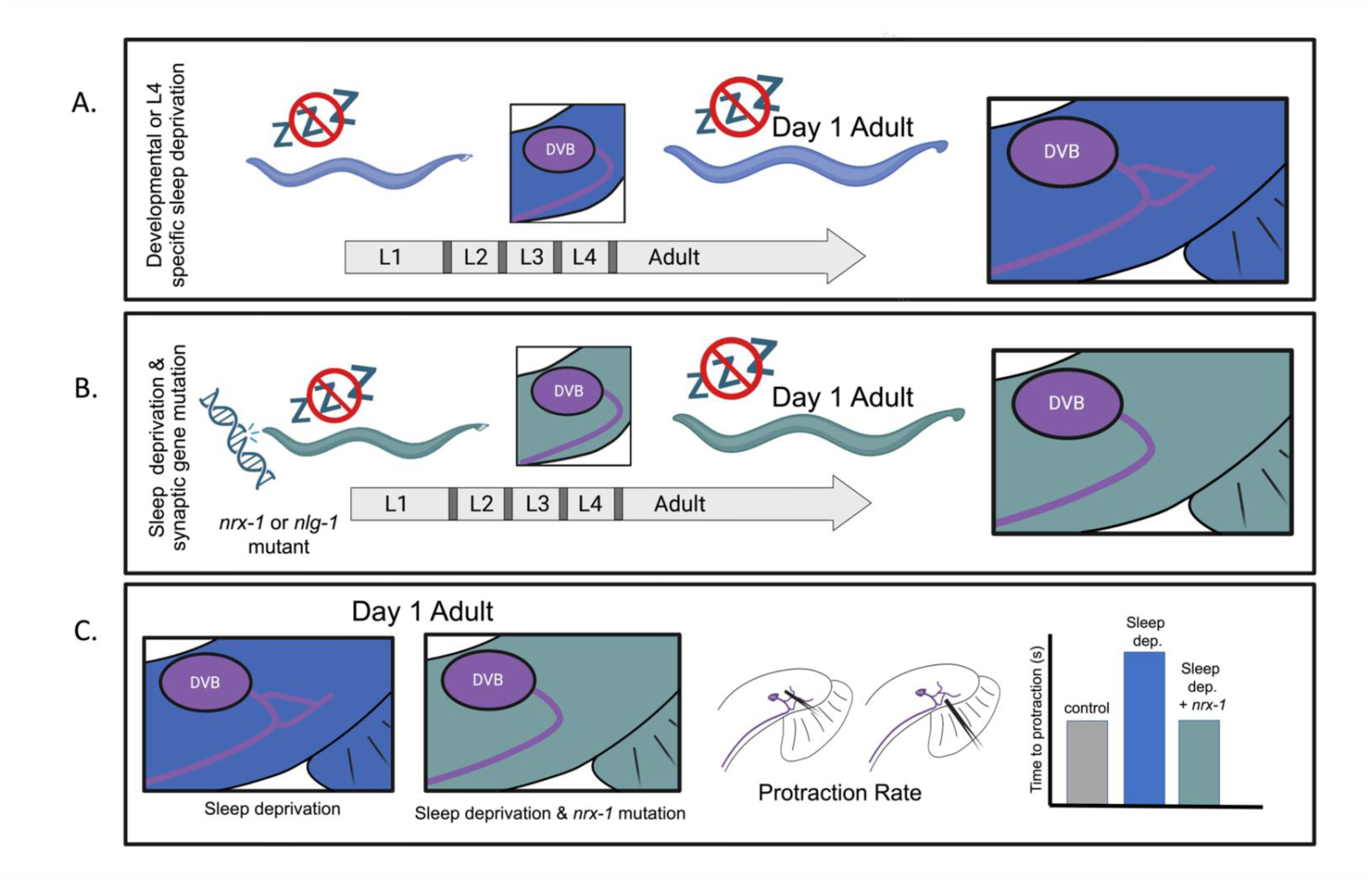
Graphical Abstract. GABAergic morphologic and behavioral plasticity induced by developmental sleep deprivation depends on the conserved autism genes, *nrx-1* and *nlg-1* **A)** Sleep depriving male *C. elegans* during all larval sleep periods in development (*aptf-1* or *npr-1*) or selectively inhibiting sleep during L4-adult molt (vibration or RIS silencing) results in an increase in DVB outgrowth. **B)** Mutations in *nrx-1* (and *nlg-1* in *npr-1* animals) prevent sleep deprivation induced morphologic plasticity changes. **C)** DVB neurite outgrowth results in a functional change in spicule protraction rate as a result of increased GABAergic inhibitory signaling, which also depends on *nrx-1* and *nlg-1*.

## INTRODUCTION

Sleep is essential for brain function and survival across species. While the neurologic benefits of sleep are still being defined, sleep is important for memory consolidation, attention, learning, plasticity, and waste clearance/metabolic regulation for neuron preservation^1–3^. Long-term sleep deprivation is associated with severe cognitive impairment^4, 5^. Additionally, sleep disruption exacerbates a number of physiological problems including neurodegenerative diseases and impedes neuronal rehabilitation after injury including stroke or traumatic brain injury^6–10^. Further, sleep deficits are linked to neuropsychiatric and neurodevelopmental conditions^11–15^ including autism, in which sleep impairments impact core behavioral characteristics^16–24^.

Structural plasticity, or the ability of neurons to reorganize their shape and connections in response to environment and experience, appears intimately tied to sleep^11, 13, 25–28^. Analysis in *C. elegans, Drosophila*, zebrafish, and rodents demonstrates that sleep regulates synapse strength, number, composition, and morphology, as well as learning, memory, mood, and social behaviors^29–45^. However, the functional role of the synaptic changes on behaviors, and the mechanisms that bridge them remain elusive. Neuronal activity during wakefulness correlates with sleep behavior itself^46^, uncovering a reciprocal relationship between sleep and plasticity. Many methods used for sleep disruption across models may also impact neuronal and circuit activity unrelated to sleep (ex. gentle handling and novel object exposure involve stress and learning, respectively)^47^. Additionally, disparate or contradictory findings in the number and strength of synapses after sleep and sleep disruption likely reflect differences in the brain region, organism, and sleep disruption method used^26, 48^. The majority of studies on sleep deprivation analyze either synaptic plasticity or behavior, but not both^30–34, 36, 37, 41, 42^. Analyses of synapses and behavior together have mainly focused on the impact of sleep deprivation on learning-induced plasticity^38, 43–45^. The behavioral effects of specific synaptic changes in response to sleep loss beyond learning induction have been difficult to determine as the neurons and circuits that regulate distinct behaviors are still being defined. Leveraging the power of *C. elegans*, we can quantify how sleep deprivation at specific time windows impacts a specific neuron (DVB) and its behavioral output (spicule protraction).

One mechanism by which sleep may modify neuron and synapse structure, resulting in behavioral changes, is through synaptic adhesion molecules (SAMs). SAMs impact synapse formation and function, mediating neuron and synaptic structural changes in development and maintenance in adults. Several adhesion molecules (immunoglobulins (IG), cadherins, neurexins, neuroligins, integrins, and Eph/Ephrins) have roles in the regulation of sleep behavior^49–53^. Neurexins are autism-associated SAM genes^54–56^ with diverse roles in synapse function and plasticity, making them particularly relevant in the context of sleep and structural plasticity^56, 57^. Alternatively spliced isoforms of neurexins can have discrete regulatory functions which are neuron-, synapse-, and temporal-specific^56–62^. While there are three mammalian neurexin genes, there is only one in *Drosophila* and *C. elegans*, thereby simplifying genetic manipulations in these model systems. Deletions in *Drosophila* neurexin (*Nrx-1*) fragment sleep behavior and disrupt circadian patterns and adult specific over-expression can improve most sleep phenotypes^30^. Mice lacking neuroligin (*Nlg1*), the canonical binding partner of neurexins, have reduced wakefulness and increased non-rapid eye movement sleep^49^. Further, sleep deprivation results in altered neuroligin and neurexin expression levels, including isoform-specific impacts^51^. The role of SAMs in sleep has relied on whole brain or brain region analysis^49, 53, 56, 63^, while work on these genes beyond sleep highlights their neuron, synapse, and context specific functions. Despite the connections between sleep, SAMs, and structural neuroplasticity, how these changes are executed and the role of SAMs in sleep deprivation-related behaviors are not well-characterized.

### Structural Plasticity of the GABAergic DVB neuron in adult male *C. elegans*

The GABAergic DVB neuron undergoes experience-dependent structural plasticity and circuit rewiring in adult male *C. elegans.* Increased DVB outgrowth, which occurs only in males, leads to changes in spicule protraction, a specific step of male mating behavior^64^. The progressive outgrowth/branching of axon-like neurites of the DVB neuron is unlike almost any other neuron in *C. elegans* in its dynamic nature and variability. DVB structural plasticity is modified by experience, such as exposure to mates or manipulation of circuit activity (stimulation or inhibition with opto-/chemo-genetics)^64^. Interestingly, DVB outgrowth is dependent on the *C. elegans* ortholog of neurexins and neuroligins, *nrx-1* and *nlg-1*. However, contrary to the canonical roles of *nrx-1* and *nlg-1* in many contexts, the two have antagonistic effects on DVB morphologic plasticity in adults, with NRX-1 promoting and NLG-1 restricting DVB neurite outgrowth^64^.

Stressors applied during an adolescent-like developmental stage (L4) lead to early DVB neurite outgrowth at day 1 of adulthood^65^. While some of these stressors (starvation) result in changes in spicule protraction behavior, other stressors (UV and heat-shock) do not impact behavior despite inducing DVB morphologic plasticity^65^. Additionally, while starvation induced DVB neurite extension is *nrx-1* and *nlg-1* dependent, neither UV nor heat-shock induced structural plasticity depend on *nrx-1* and only heat-shock depends on *nlg-1*. Despite their antagonistic functions in adulthood, *nrx-1* and *nlg-1* both promote adolescent starvation stress induced DVB plasticity at day 1^65^. Starvation stress coincides with the fourth larval stage and molt when *C. elegans* undergo a period of developmental sleep, leaving open the question of whether sleep may play a role in DVB plasticity^65^.

### C. elegans Sleep

Studying sleep in *C. elegans* is particularly advantageous given its transparency, neuronal gene conservation, and mapped 300-neuron connectome^66–68^. Further, many cell– and gene-specific tools for sleep manipulation and technologies for measuring sleep have been developed in *C. elegans*^69–78^. *C. elegans* undergo periods of behavioral quiescence during the four larval transitions, known as lethargus or developmentally timed sleep, that include all of the characteristics of sleep (cessation of movement, rapid reversibility, decreased responsiveness, stereotypical posture, reduced neuronal activity, and homeostatic regulation)^79–83^. Developmental sleep is regulated by the conserved clock Period gene^82^ and is promoted by the RIS neuron^84–87^. Disruption of APTF-1 or LIM– 6, transcription factors required to specify the RIS neuron, results in decreased sleep^88–90^. Sleep in *C. elegans* has also been shown to coincide with synaptic plasticity^91^ and recent work has defined the neuronal and synaptic changes that link experience and sleep to memory-related behavior^29^.

Here we use multiple methods to disrupt developmental sleep in *C. elegans*, including genetic, physical, and chemo-genetic manipulations. We find that sleep deprivation increases neurite outgrowth of the DVB GABAergic neuron and modifies behavior in early adulthood. We show that sleep deprivation during the adolescent-like L4 molt induces similar morphologic and behavioral changes to sleep deprivation throughout development, narrowing the critical timing of sleep to the last larval stage. Importantly, sleep deprivation induced morphologic and behavioral plasticity depend on the conserved SAMs, neurexin/*nrx-1* and neuroligin/*nlg-1*, which act in the same pathway to promote DVB structural plasticity after sleep loss. These findings show that sleep is essential for proper maintenance of neuronal morphology at single neuron and behavior resolution. Additionally, our results indicate that sleep alters behavior by modifying plasticity at the level of specific synaptic and circuit connections. This work implicates neurexins and neuroligins in sleep dependent structural plasticity and identifies isoform specificity of *nrx-1*. Taken together, our results show that sleep disruption robustly alters DVB structural plasticity and behavior, and these changes are dependent on conserved SAMs that are associated with autism and other neurologic conditions.

## METHODS

### *C. elegans* strain maintenance

All strains were maintained on NGM-agar filled plates and seeded with OP50 bacteria as a food source. All strains tested had a *him-8* mutation to increase the proportion of male *C. elegans.* Experiments were performed only on males since DVB adult structural plasticity occurs only in males^64^. Transgenic worms used in this study were previously published and either existed in the lab or were provided by others^65, 92^. Strains, mutant alleles, and transgenics included are listed in **Table 1** by figure order.

### Confocal microscopy

For day 1 experiments, *C. elegans* were picked onto new plates at larval stage 4 based on morphology and size, then subjected to confocal imaging the following day (day 1 adult) or three days later (day 3 adults). Males were imaged on a Leica SP8 point scanning confocal microscope using a 63x objective with glycerol immersion fluid (type G). Worms were mounted on an agar pad made of 5% agarose and paralyzed in a 100mM sodium azide solution in M9. The DVB neuron was imaged at 2x zoom. Z-stacks were set such that imaging spanned the entire neuron at a slice size of 0.6μM. Z-stacks were approximately 30-75 slices. Representative image figures were made using FIJI image J and Adobe Illustrator.

### DVB neuron morphology analysis

DVB neuron morphology analysis was performed in FIJI image J. Analysis was performed using the Simple Neurite Tracer or updated SNT Neuroanatomy plugin. Neurites were traced as previously described ^64, 65^. Briefly, skeletons were created starting at the center of the DVB soma and extending to the longest point of neurite extension through the Z– stacks. Branches were then traced with the start site connected to the existing trace. All posteriorly extending branches of DVB were traced to the anterior turn of DVB on the ventral side. Total neurite length was determined using the Analyze Skeleton function. Data were graphed in GraphPad Prism 9.

### DVB *cla-1::gfp* pre-synaptic puncta analysis

Puncta in control, *aptf-1*, and *npr-1* mutant day 1 males were analyzed using the FIJI Analyze Particle function, which allows for unbiased quantification of *cla-1::gfp* puncta. *lim-6int4p::cla-1::gfp* transgenic worms were crossed with the otIs541 (*lim*– *6int4p::wCherry*) strain and imaged at 63X on the Leica SP8 point scanning Confocal Microscope with an additional zoom of 2.5x. Z-stacks were set as described above. Lightning was applied to enhance resolution. Lightning deconvolution applies an appropriate reconstruction strategy to individual voxels rather than using a single parameter on the whole image to extract detailed image information^93^. Micrographs were converted to 8-bit and auto-thresholded. If auto-thresholding was not representative of *cla-1::gfp* puncta, minor manual adjustment was made to match original images most closely. A region of interest (ROI) was defined to restrict puncta counts to DVB neurites (exclusion of soma and PVT neuron). Analyze Particles generated *cla-1::gfp* counts and bare outline images for the ROI. Pixel size of *cla-1::gfp* puncta was restrained to 0.01 pixels and circularity was not restricted. Data were graphed in GraphPad Prism 9. Traces for representative images were made in Adobe Illustrator and bare outlines of *cla-1::gfp* puncta were super-imposed on traces.

### Vibration Sleep Deprivation

Mid Larval stage 4 males, which were identified based on morphology, were transferred to new NGM plates coated with OP50 bacteria. Plates housing the worms were then placed inside the WormWatcher platform developed by Tau Scientifics^94^. One-second– long 1Hz vibration was applied every ten seconds for a duration of six hours across the L4 molt. Adults were imaged for DVB morphology as described above. For vibration experiments before the L4 molt, worms picked as L3s based on morphology and stimulus was applied four hours later. For vibration experiments after the L4 molt, worms were picked as early L4s and vibration was applied ten hours later when the animals were early adults.

### Histamine silencing of RIS neurons

To silence RIS, transgenic worms expressing a histamine gated chloride channel in RIS (NQ1208, *flp-11p::HisCl::SL2::mCherry)* were used^92^. Males with this transgene were identified based on expression of the co-injection marker *myo-2p::mCherry* and picked at larval stage 4. L4 *C. elegans* were placed on top of an individual NGM-filled well of a WorMotel^76, 94^,. 1.5μL of 10mM histamine in water was applied to the surface of WorMotel wells filled with NGM agar. Following an incubation period of one hour to allow the histamine to diffuse into the agar, L4 males were transferred to the WorMotel with food (OP50) spread on each well. WorMotels were imaged using WormWatcher Platforms (Tau Scientific) with a picture taken every ten seconds for ten hours. A MATLAB GUI^76^ was used to apply pixel subtraction between images to determine activity and quiescence of worms. Day 1 adults were subjected to confocal microscopy to measure DVB morphology as described above.

### Aldicarb spicule protraction assay

Larval stage 4 males of each tested genotype/condition were transferred to a new NGM plate coated with OP50. 8-15 worms were placed on each plate. For vibration sleep deprivation, worms were placed in a WormWatcher and subjected to the above vibration protocol. Males were tested the following day (day 1 adults) on the previously described aldicarb spicule protraction assay^64^. One hour prior to experiments, 130μL of 100mM aldicarb were applied to empty NGM plate, without OP50, and evenly spread across the plate. Aldicarb was allowed to absorb into the plate and dry for one hour. Following transfer to the aldicarb assay plate, we measured the latency to spicule protraction lasting at least five seconds. Males were removed from the plate after spicule protraction. Time to protraction was plotted in GraphPad Prism 9.

### Statistical analysis

All data were plotted and analyzed in GraphPad Prism 9. To determine statistical significance, we used one-way ANOVA with Tukey’s post-hoc test. For comparisons of two data sets, a two-tailed unpaired t-test was used to compare significance. Number of worms used were based on previous studies and effect size. “n” represents the number of individual worms tested and is indicated in each figure. Each morphology or behavior experiment was performed at least 3 times, with each trial performed on a different day. Error bars on figures represent standard error of the mean (SEM). P-values are shown in each figure.

## RESULTS

### Developmental sleep disruption increases DVB neurite outgrowth in a *nrx-1* dependent manner

To determine whether sleep deprivation alters structural plasticity, we studied neuron morphology of the sexually dimorphic GABAergic neuron, DVB, in males at day 1 of adulthood. We compared control strains, in which animals slept normally to two strains with disrupted sleep across development (*aptf-1(tm3287)* and *npr-1(ad609)* mutants)**(Figure 1A)**^90, 95^. Worms lacking APTF-1, a conserved AP2 transcription factor, have profoundly decreased sleep at all periods of developmental sleep as APTF1 is a positive regulator of FLP-11, a sleep-inducing neuropeptide^88, 90^. The role of AP2 transcription factors in sleep regulation is conserved in *Drosophila*, mice, and humans^96–98^. We found an increase in DVB neurite length in *aptf-1(tm3287)* mutants with disrupted sleep, on day 1 of adulthood **(Figure 1C&E)**. Additionally, day 1 adult *aptf-1(tm3287)* males have an increase in the number of DVB junctions, a proxy for neurite branching compared to controls **(Figure 1D&E)**. NPR-1 encodes a neuropeptide Y like receptor and mutants lacking NPR-1 have decreased sleep due to increased sensory activity, mediated by the arousal neuropeptide PDF-1^85, 95, 99^. DVB neurite length was significantly increased in day 1 adult *npr-1(ad609)* mutant males compared to controls **(Figure 1F-H)**.

**Figure 1:**
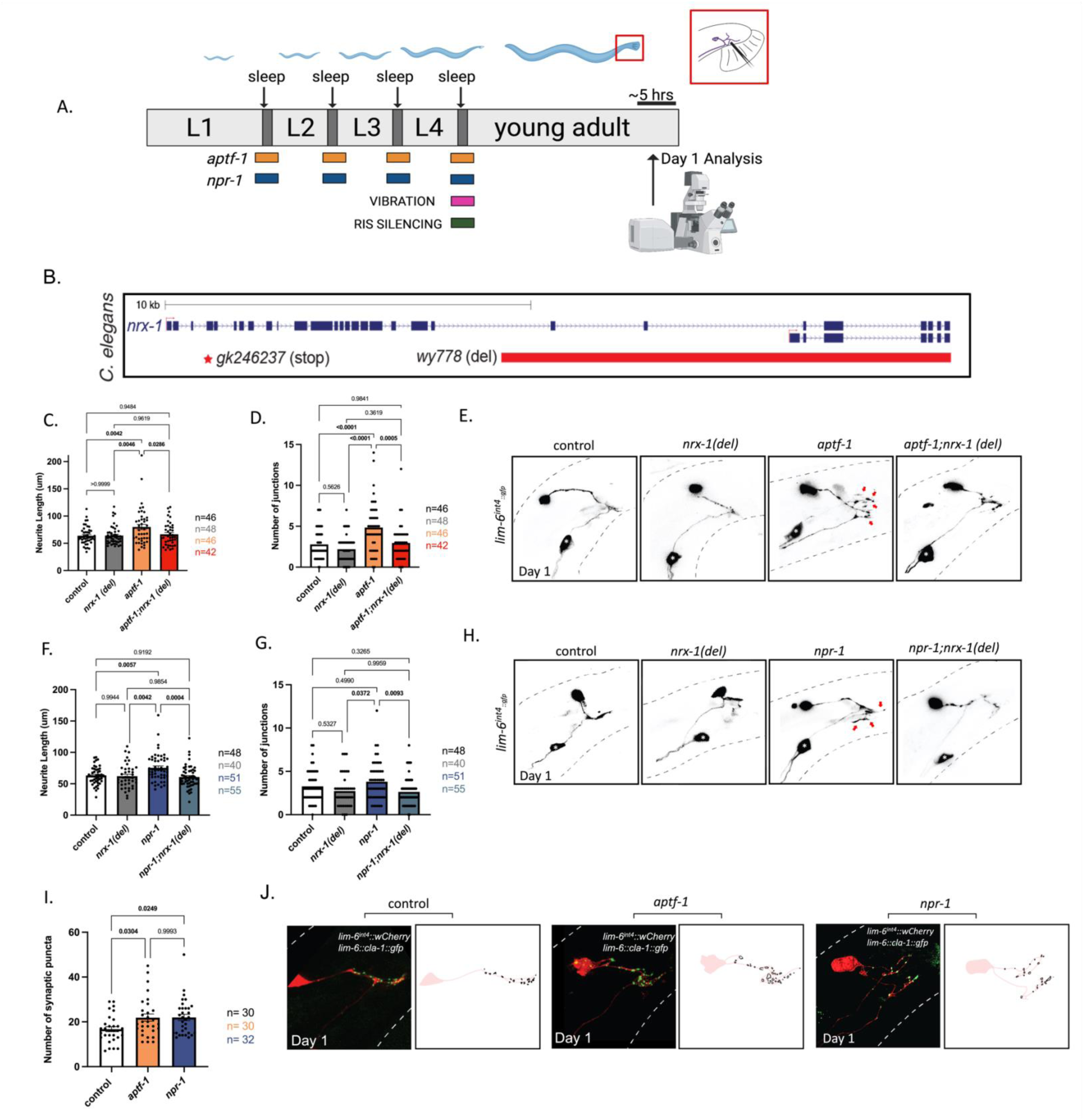
Genetic disruption of developmental sleep (via *npr-1* and *aptf-1*) increases DVB neurite outgrowth dependent on *nrx-1*. **A**) Schematic of *C. elegans* developmental sleep stages between larval stages with timing of each sleep deprivation method indicated. *aptf-1* (orange) and *npr-1* (blue) animals experience impaired sleep during all larval sleep periods, while vibration (pink) and RIS silencing (green) are specific to L4 to adult transition/molt. Cartoon zoom in of the adult male tail with DVB shown. **B)** Schematic of *C. elegans nrx-1* gene coding locus, red box indicates deletion in *nrx-1(wy778)* referred to as *nrx-1(del)*, red star indicates premature stop in *nrx-1(gk246237)* referred to as *nrx-1(stop).* Quantification of DVB **(C)** total neurite length and **(D)** junctions in controls, *nrx-1(del), aptf-1*, and *aptf-1; nrx-1(del)* males at day 1. **E**) Representative images of DVB neuron in day 1 males using *lim-6p::gfp.* Red arrows indicate neurite branches. Black dashed lines represent body outline. White asterisk marks PVT neuron (not traced, see^64^). Quantification of DVB **(F)** total neurite length and **(G)** junctions in controls, *nrx-1(del), npr-1*, and *npr-1; nrx-1(del)* day 1 males. **H)** Representative images of DVB at day 1 in *npr-1* mutant males and all controls. **I)** Quantification of number of *cla-1::gfp* puncta in DVB in controls and sleep deprived *aptf*– *1* and *npr-1* males at day 1. **J)** Representative images of DVB (red) and *cla-1::gfp* puncta (green) in controls, *aptf-1*, and *npr-1* day 1 males. Outlines of synaptic puncta generated from FIJI particle analysis are shown (black) superimposed on a trace of the DVB neuron (light red). Number of individual animals is indicated by “n”, P-values from one-way ANOVA with Tukey’s post-hoc test shown.

To determine whether neurexin/*nrx-1* plays a role in sleep deprivation induced neurite extension, we tested a *C. elegans* strain carrying a *nrx-1* mutation *(wy778)*, a ∼12,000 nucleotide deletion, that disrupts the long (alpha) and short (gamma) isoforms, referred to as *nrx-1(del)* in figures **(Figure 1B)**. The increase in DVB neurite outgrowth observed in *aptf-1(tm3287)* males at day 1 is lost in *aptf-1(tm3287); nrx-1(wy778)* males **(Figure 1C-E)**. Similarly, the increased DVB neurite outgrowth we observed in *npr-1 (ad609)* mutants was eliminated in *npr-1(ad609); nrx-1(wy778)* double mutants **(Figure 1F-H).** *nrx-1(wy778)* alone does not impact either measure of DVB neurite outgrowth **(Figure 1C,D,F,G)** indicating that *nrx-1* is not needed for day 1 DVB morphology, but rather, is required for sleep deprivation-induced DVB neurite outgrowth, similar to previously tested stressors^64, 65^.

We next asked whether DVB neurite outgrowth induced by sleep deprivation in male *C. elegans* corresponds to any changes in DVB pre-synaptic morphology. We expressed GFP tagged active zone marker, *cla-1*, in DVB using the *lim-6* promoter^65, 100^. Day 1 *aptf-1(tm3287)* and *npr-1(ad609)* males have an increased number of DVB *cla-1* puncta compared to controls **(Figure 1 I&J)**. The size of *cla-1* puncta was not different in sleep deprived males compared to controls **(Supplemental Figure 1)**. Therefore, sleep deprivation results in increased pre-synaptic puncta in the DVB neuron.

### Adolescent specific sleep deprivation induces DVB neurite outgrowth

Our previous work implicated the adolescent-like stage L4, when *C. elegans* males sexually mature, as particularly sensitive for induction of DVB plasticity by stress^65^. To ask if sleep deprivation during the L4 larval sleep period impacts DVB outgrowth in early adulthood, we applied a vibration stimulus during the L4 molt and analyzed DVB morphology in day 1 adult males **(Figure 1A)**. Vibration was applied for one second every ten seconds during a six-hour period overlapping with the L4 molt sleep period. L4 vibration increased DVB neurite outgrowth and number of junctions relative to controls **(Figure 2A-C)**. Vibration may impact DVB outgrowth unrelated to sleep through stimulation of sensory neurons or other stimulation of stress^101^. However, we found that vibration only impacts DVB neurite outgrowth when applied during L4 developmental sleep; the same vibration protocol applied before or after the L4 sleep period had no impact on DVB morphology **(Supplemental Figure 2)**. This result demonstrates that the L4 sleep period is specifically critical for the regulation of DVB structural plasticity.

**Figure 2:**
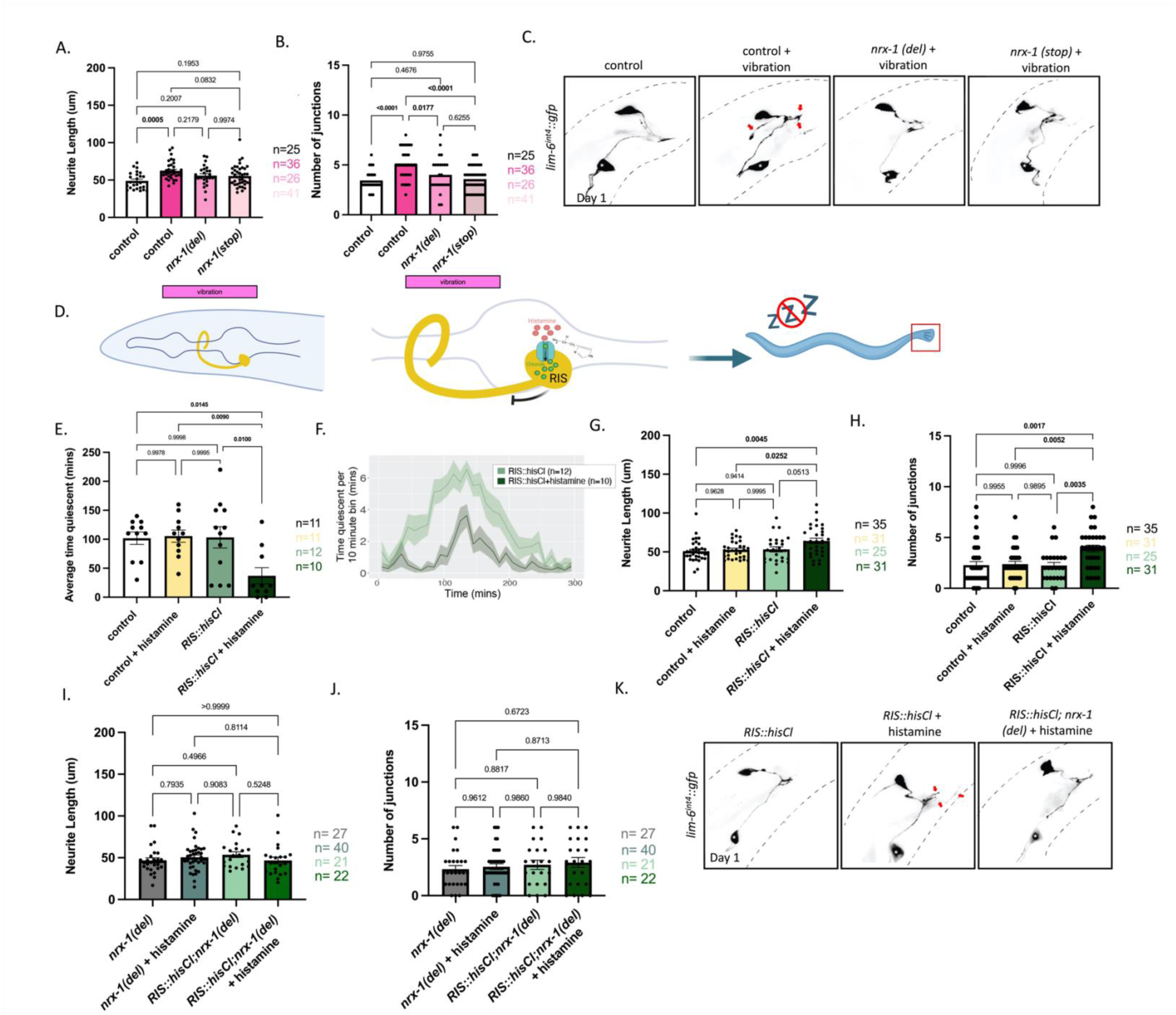
Sleep deprivation specifically during adolescent-like stage (L4) induces DVB neurite outgrowth at day 1. Quantification of DVB **(A)** neurite length and **(B)** number of DVB junctions in controls, vibrated controls, and vibrated *nrx-1* (del), and vibrated *nrx-1 (stop)* males at day 1 after vibration sleep deprivation (1Hz vibration during L4 molt – 1 second every 10 seconds for 6 hours). Vibration sleep deprivation protocol indicated with pink boxes. **C)** Representative images of DVB neuron in vibration sleep deprived males and controls. White asterisk marks PVT neuron (not traced, see^64^) **D).** Cartoon of histamine gated chloride channel expression in RIS neuron to silence RIS neuron activity. Exogenous histamine acts to open chloride channels allowing for the influx of negative ions to hyper-polarize the neuron and prevent signaling, selectively in the RIS neuron (*flp-11* promoter), referred to as *RIS::hisCl.* **E)** Graph of time quiescent during L4 molt in *RIS::hisCl* transgene positive and negative males with (dark green) and without (light green) histamine application. **F)** Aligned quiescence peak during L4 lethargus/molt in *RIS::hisCl* transgenic male *C. elegans* with and without (+/−) histamine application during L4 to adult transition. Amount of sleep is binned in 10-minute segments. **(G)** DVB neurite length and **(H)** junctions in *RIS::hisCl* transgenic day 1 males +/− histamine and controls. **I)** DVB neurite length and **(J)** junctions in *RIS::hisCl; nrx-1(del)* +/− histamine and *nrx-1(del)* +/− histamine day 1 males. **K)** Representative images of DVB neuron in *RIS::hisCl* transgenic +/− histamine and *RIS::hisCl; nrx-1(del)* day 1 males. White asterisk marks PVT neuron (not traced, see^64^). Number of individual animals is indicated by “n”, P-values from one-way ANOVA with Tukey’s post-hoc test shown.

To disrupt sleep in another fashion, we next silenced the sleep promoting neuron RIS through expression of a histamine-gated chloride channel (*flp-11* promoter)^92^. Application of histamine results in hyperpolarization of the RIS neuron, preventing it from releasing GABA and the sleep-promoting neuropeptide FLP-11, and allowing temporal control of sleep loss **(Figure 2D)**. Using a WorMotel^76, 94^, we monitored the impact of histamine in transgenic and non-transgenic animals on quiescence and activity for 10 hours to capture the L4 sleep period (5 hours shown around and normalized to peak quiescent hour) **(Figure 2F).** Overnight histamine exposure during the L4 sleep period in males expressing *flp-11p::HisCl* significantly reduced quiescence, a proxy for sleep quantification on the WorMotel^76^, compared to *flp-11p::HisCl* males without histamine (37 minutes vs. 105.5 minutes) **(Figure 2E&F)**. Histamine on its own did not impact quiescence. Thus, we can disrupt sleep specifically and temporally using this chemo– genetic approach.

Male *C. elegans* with sleep disruption via histamine at L4 displayed increased adult DVB neurite outgrowth compared to controls. We observed longer day 1 neurite lengths in L4– stage RIS-silenced animals (64.08uM) compared to untreated control animals (53.24 μM)(P=0.051, **Figure 2G&K**). RIS-silenced males had significantly increased DVB neurite length compared to control animals not carrying the HisCl transgene (N2 on histamine, N2 without histamine)**(Figure 2G&K)**. RIS-silenced day 1 adult males had significantly increased DVB junctions compared to HisCl transgenic animals untreated with histamine and to controls not carrying the HisCl transgene, either treated or untreated with histamine **(Figure 2H&K)**. This result confirms the importance of the L4 sleep period in regulation of DVB structural plasticity.

*nrx-1* can have isoform-specific functions^102^ so we asked whether DVB outgrowth after sleep disruption was dependent on any specific isoform of *nrx-1*. We compared the *nrx-1*(*wy778)* allele used above with *nrx-1 (gk246237)*, a premature stop allele that specifically disrupts the alpha (long) isoform, referred to as *nrx-1(stop)* in figures **(Figure 1B)**. Vibration based sleep deprivation did not increase DVB neurite length in either mutant allele of *nrx-1* compared to controls **(Figure 2A&C)**. The number of DVB junctions was significantly increased in vibration sleep deprived males compared to *nrx-1(wy778)* and *nrx-1 (gk246237)* vibration sleep deprived males **(Figure 2A-C)**. This result suggests that the long isoform of *nrx-1* specifically is required for DVB neurite outgrowth resulting from vibration sleep disruption.

We next tested *nrx-1(wy778)* mutants with the *flp-11p::HisCl* transgene in the presence or absence of histamine application. In alignment with our other findings, RIS– silenced *nrx-1(wy778)* mutants do not have increased neurite length or junctions relative to non-transgenic *nrx-1(wy778)* mutants on histamine **(Figure 2I-K)**. This result confirms and extends our results from genetic and vibration sleep deprivation that *nrx-1* regulates the morphological changes caused by L4 sleep deprivation.

### Sleep deprivation induced DVB structural plasticity alters behavior

Increased DVB neurite outgrowth in adulthood leads to changes in spicule protraction behavior, which is the readout of the balance between excitation from cholinergic SPC neurons and inhibition from the GABAergic DVB neuron **(Figure 3A)**^103^. We used the previously described assay in which spicule protraction is induced by application of aldicarb, an acetylcholinesterase inhibitor that blocks breakdown of acetylcholine and promotes spicule protraction^64^. We monitored spicule protraction at day 1 on aldicarb **(Figure 3 A&B)**^64^ and found that *aptf-1(tm3287)* males took longer to protract their spicules compared to controls **(Figure 3C)**. Day 1 *npr-1(ad609)* males also showed an increase in time to spicule protraction relative to controls **(Figure 3D)**. As was the case with *aptf-1* and *npr-1* sleep-deficient mutants, we found that sleep disruption at the L4 molt via vibration significantly increased time to spicule protraction at day 1 **(Figure 3E)**. Previously, we showed that *nrx-1(wy778)* day 1 males do not alter time to spicule protraction^65^.

**Figure 3:**
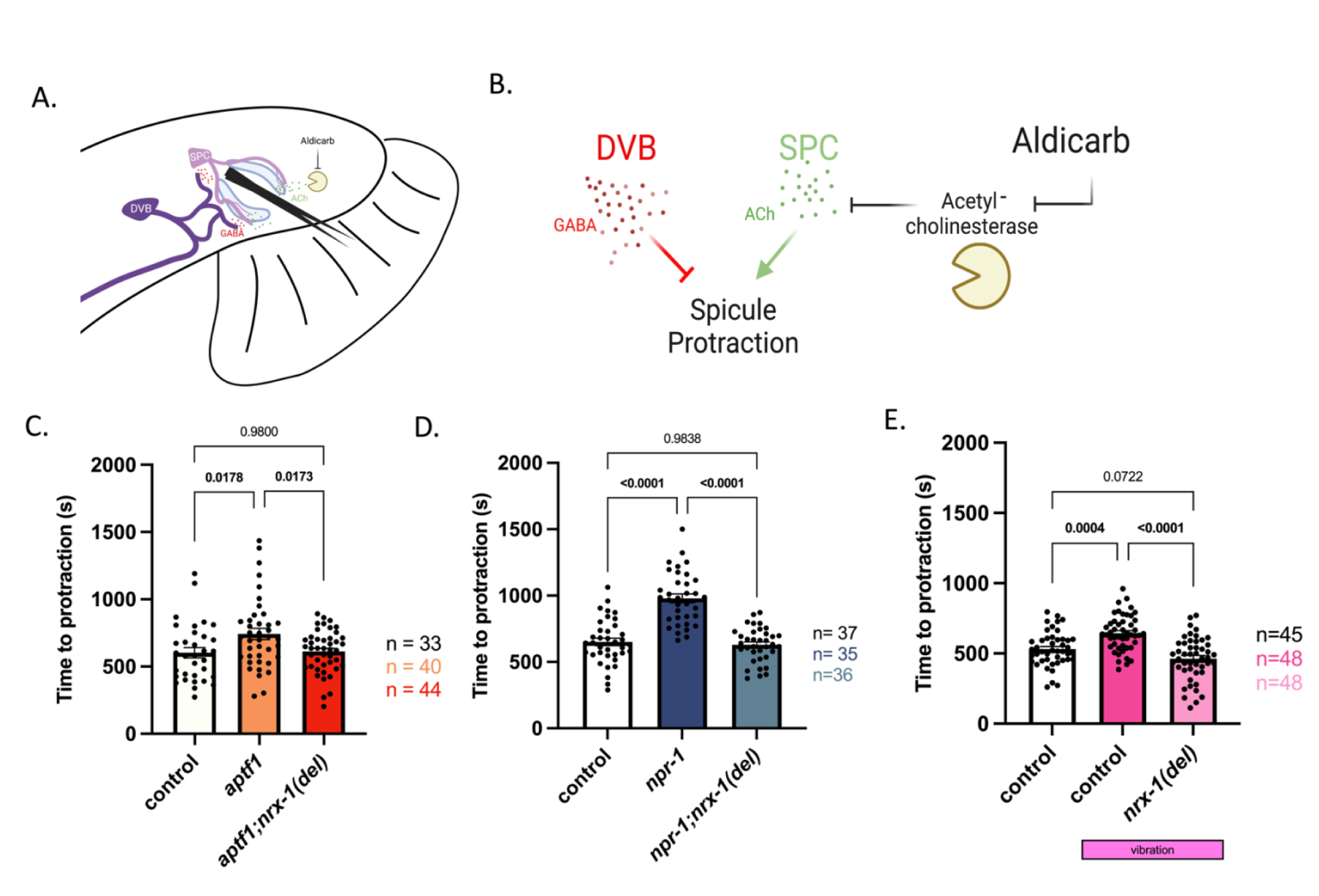
Sleep deprivation alters DVB-dependent spicule protraction behavior. **A**) Cartoon of neurons and muscles controlling spicule protraction in male tail. DVB is shown in dark purple and SPC is shown in light purple. **B)** Cartoon depicting mechanism for aldicarb induced spicule protraction with opposing functions of DVB GABAergic inhibition (red) and SPC cholinergic excitation (green). **C)** Time to spicule protraction following aldicarb exposure in control, *aptf-1*, and *aptf-1; nrx-1(del)* day 1 males. **(D)** Time to spicule protraction following aldicarb exposure in control, *npr-1*, and *npr-1; nrx-1(del)* day 1 males **E).** Time to spicule protraction following aldicarb exposure in control day 1 males and control and *nrx-1(del)* day 1 males that underwent L4 vibration sleep deprivation. Number of individual animals is indicated by “n”, P-values from one-way ANOVA with Tukey’s post-hoc test shown.

To determine any role for *nrx-1* in behavioral changes following sleep deprivation we tested *aptf-1(tm3286); nrx-1(wy778)* and *npr-1(ad609); nrx-1(wy778)* double mutants. We found that the increased time to spicule protraction in the sleep mutants and after L4 vibration is suppressed by loss of *nrx-1* **(Figure 3C-E)**. These results show that sleep deprivation impacts DVB-dependent behavior and indicate that neurite outgrowth induced by sleep loss has functional consequences. Further, we show that neurexin/*nrx-1*, which is required for sleep deprivation induced morphologic plasticity, is also required for behavioral plasticity.

### DVB neurite outgrowth and behavior after sleep deprivation is dependent on *nlg-1*

To determine whether neurexin’s canonical binding partner, neuroligin/*nlg-1*, also contributes to sleep deprivation induced DVB plasticity and behavior, we tested the impact of a large deletion allele, *nlg-1(ok259)* **(Figure 4A).** We found that increased DVB neurite length in *npr-1* mutant males was suppressed by *nlg-1(ok259)* **(Figure 4B-D)**. *npr*– *1(ad609); nlg-1(ok259)* males and *nlg-1(ok259)* mutants alone were not significantly different from controls **(Figure 4B-D)**. *nlg-1(ok259)* mutation also suppressed the increased latency to spicule protraction observed in *npr-1* single mutants **(Figure 4E).** These results demonstrate that *nlg-1, like nrx-1*, regulates both DVB structure and function in response to sleep deprivation.

**Figure 4:**
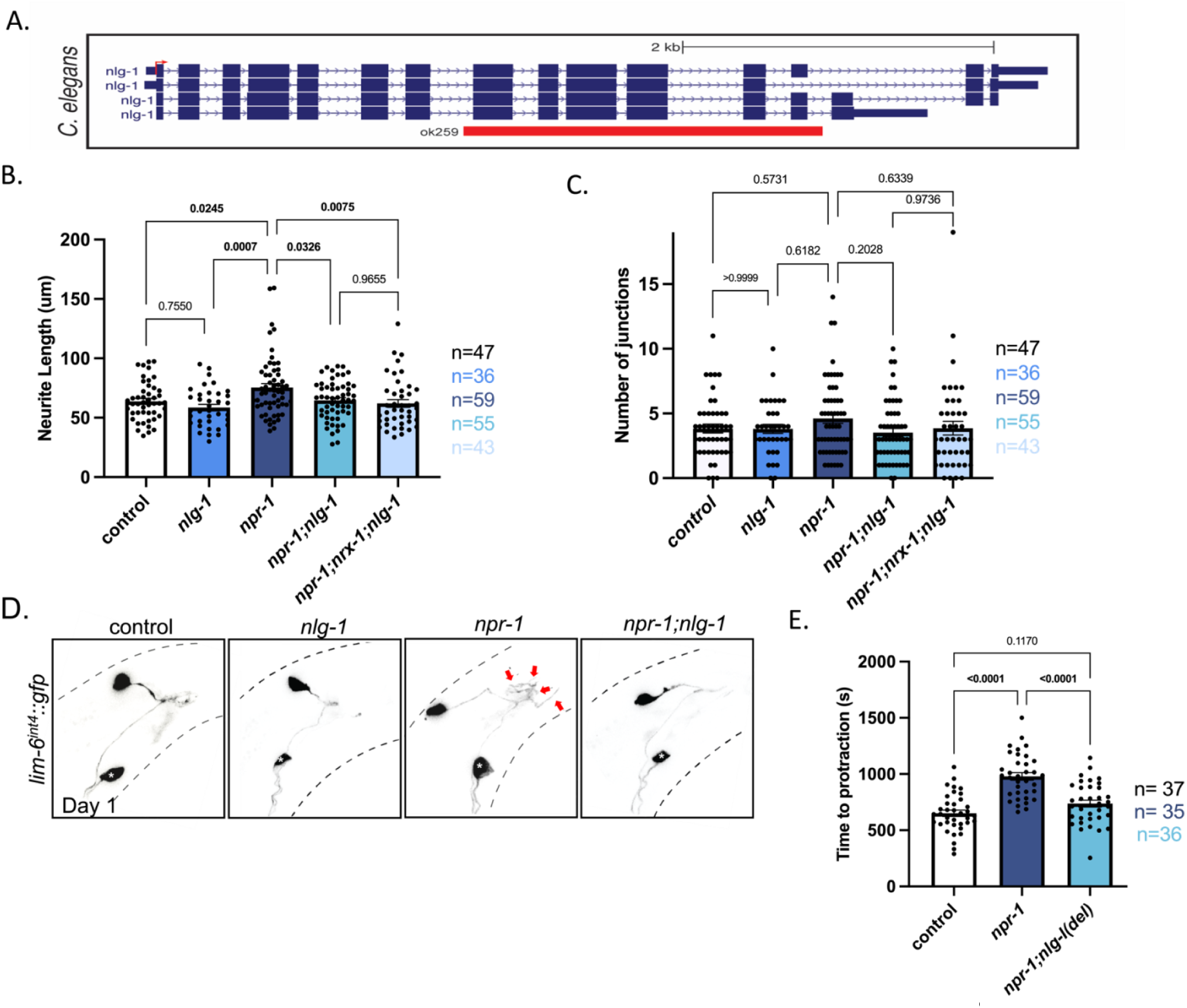
*nlg-1* is required for sleep deprivation induced DVB neurite outgrowth and spicule protraction behavioral changes. **A)** Schematic showing *nlg-1 C. elegans* gene locus with *nlg-1(ok259)* deletion indicated with red box. **B)** Graph displaying quantification of DVB neurite length and **(C)** DVB junctions in controls, *nlg-1, npr-1, npr-1; nlg-1*, and *npr-1; nrx*– *1; nlg-1* day 1 males. **D)** Representative images of DVB neuron in controls, *nlg-1, npr-1*, and *npr-1; nlg-1* day 1 males. White asterisk marks PVT neuron (not traced, see^64^). **E)** Time to spicule protraction following aldicarb application of control, *npr-1*, and *npr-1; nlg-1* day 1 males. Number of individual animals is indicated by “n”, P-values from one-way ANOVA with Tukey’s post-hoc test shown.

*nrx-1* and *nlg-1* may act in concert to control sleep deprivation induced plasticity of DVB in which case there would be no further suppression of the sleep deprivation induced phenotypes in animals lacking both *nrx-1* and *nlg-1.* DVB neuronal morphology in *npr*– *1(ad609); nrx-1(wy778); nlg-1(ok259)* triple mutants did not differ from *npr-1(ad609)* mutants with single mutations in either *nrx-1* or *nlg-1* **(Figure 4B-D)**. This result suggests that *nrx-1* and *nlg-1* act in the same molecular pathway to regulate sleep deprivation induced DVB morphologic plasticity. Another possibility is that *npr-1; nrx-1; nlg-1* triple mutants do not show a further decease in neurite length due to a floor effect; however, we observe that DVB branching and length in these day 1 adults is higher than both L4 animals and hermaphrodites^64^.

### DVB neurite outgrowth after developmental sleep deprivation is transient

DVB neurite outgrowth increases from day 1 to day 5 of adulthood and adolescent stress leads to lasting changes in DVB morphology at least to day 3 of adulthood^64, 65^. We asked if DVB neurite outgrowth at day 1 after sleep deprivation persists into later days of adulthood. Disruption of developmental sleep via *aptf-1(tm3287)* or *npr-1(ad609)* mutations, which increases neurite outgrowth at day 1, had no difference in DVB neurite outgrowth when analyzed at day 3 **(Figure 5A-F, Supplemental Figure 3).** Thus, the DVB morphologic changes following developmental sleep deprivation observed at day 1 are transient and not long lasting, distinguishing sleep deprivation from other adolescent stressors, which have long-term changes in DVB neurite morphology^65^.

**Figure 5:**
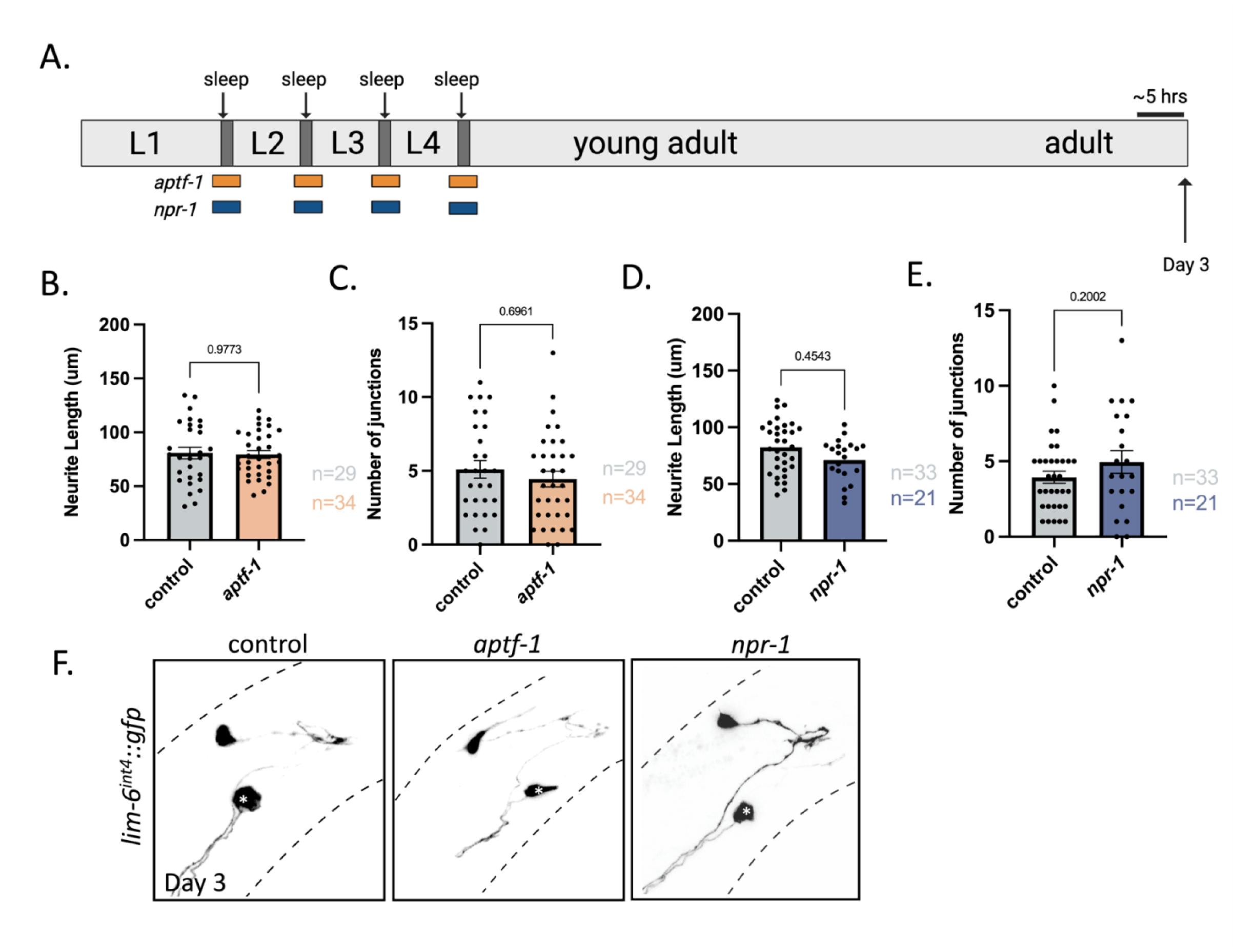
DVB neurite outgrowth induced using genetic developmental sleep disruption is transient. **A**) Schematic of experimental set-up for day 3 imaging of DVB neuron in sleep deprivation mutants, *aptf-1* (orange) and *npr-1*(blue). **B)** Graph showing DVB neurite length and **(C)** junctions in day 3 males in controls and *aptf-*1 males. **D)** Day 3 DVB neurite length and **(E)** junctions in controls and *npr-1* males. **F)** Representative images of day 3 DVB neurons in control*, aptf-1*, and *npr-1* males. The effect of *nrx-1* and *nlg-1* mutations on day 3 DVB neurite outgrowth was not tested as there was no difference between sleep deprived and control *C. elegans*. White asterisk marks PVT neuron (not traced, see^64^).Number of individual animals is indicated by “n”, P-values from two-tailed unpaired t-test shown.

## DISCUSSION

We previously described structural plasticity of the GABAergic neuron DVB in male *C. elegans* during the first 5 days of adulthood ^64^ and altered plasticity following adolescent stress^65^. Changes in DVB morphology and behavior are dependent on the single neurexin *(nrx-1)* and neuroligin *(nlg-1)* in *C. elegans*, two highly conserved synaptic adhesion molecules (SAMs) with links to autism^54–56^. How sleep, structural neuroplasticity, autism genes, and behavior influence one another requires a system in which to simultaneously manipulate and analyze each interaction. Here, we disrupted developmental and adolescent sleep in *C. elegans* and measured the impact on the morphologic and behavioral plasticity of the DVB neuron^64, 65^. We found that all four methods of sleep disruption tested (encompassing genetic, physical, and chemo-genetic manipulations) altered plasticity of the DVB neuron, resulting in changes in morphology and behavior at day 1 of adulthood. Sleep disruption specifically during larval stage 4 induced neurite outgrowth and behavioral changes, though future work will be required to test the role of other developmental sleep periods. Additionally, we find that the neuronal changes induced by sleep loss depend on *nrx-1* and *nlg-1*, acting in the same molecular pathway. In surprising contrast to UV, heat-shock, and starvation which affects DVB outgrowth at day 3 after adolescent stress^65^, sleep deprivation induced DVB morphologic changes are transient. Our results indicate that sleep (developmentally and specifically at L4 lethargus) is an important factor in regulating structural plasticity and behavior and that autism-associated gene mutations alter the neuronal response to sleep loss.

Our findings lead us to a model in which overall DVB neurite outgrowth is determined by the early balance of DVB neurite extension and retraction, as supported by our previous work which shows these dynamic changes in DVB neurites **(Figure 6A)**^64^. Selective pruning and strengthening of synapses by neurons allows for tuning of neural circuits and can impact learning ^45, 104^. During adulthood, projections to nonessential targets may form and get pruned back while projections to other targets, like SPC and spicule muscles, are stabilized or increased^64^. In turn, males become more efficient at mating, likely reflecting improved coordination of spicule protraction. In support of this, day 1 males show aberrant spicule protractions during defecation that is ameliorated by day 3 due to increased projections and inhibition by DVB^64^.

**Figure 6:**
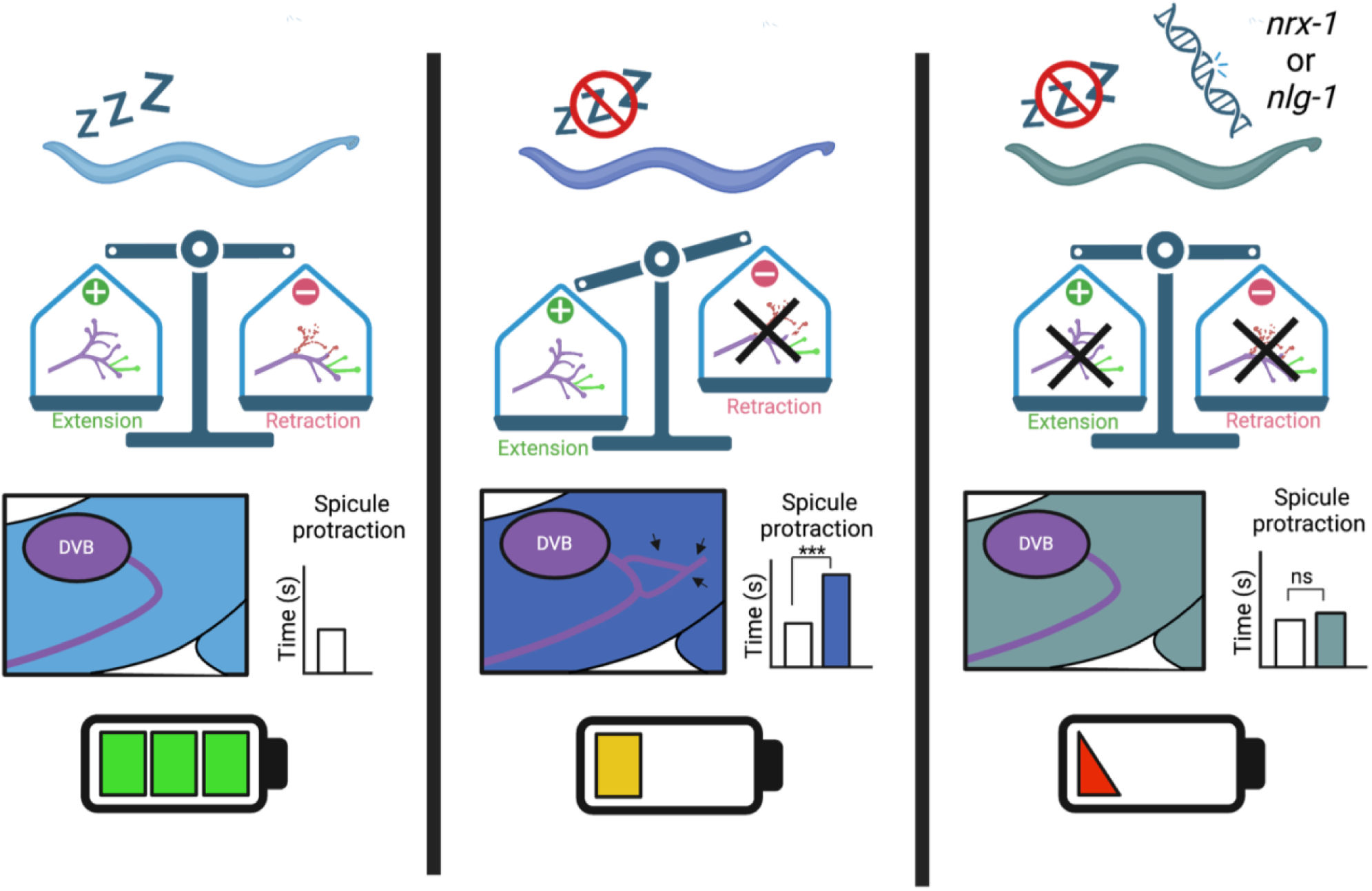
Model for sleep regulation of DVB neurite outgrowth and behavior. **A**) We propose that DVB morphology in day 1 male *C. elegans* is determined by the balance of neurite extension and retraction (or pruning) after the L4 transition to adult. DVB neurite outgrowth leads to inhibition from DVB, which determines the time to spicule protraction. *C. elegans* with undisturbed sleep have a balance in extension and retraction of neurites and conserve energy during quiescent periods. **B)** We suggest that DVB undergoes neurite extension without retraction in male *C. elegans* that experience sleep loss, with pruning being dependent on sleep, thus resulting in an imbalance between extension and retraction and an overall increase in DVB neurite outgrowth. The increase in DVB neurites results in an increase in DVB inhibition and a longer time to spicule protraction. While energy is not conserved during quiescence in these animals, the increase in inhibition on spicule neurons and muscles, observed as slower spicule protraction, may reduce energy expenditure. **C)** In day 1 animals lacking sleep with mutations in *nrx-1* or *nlg-1*, we hypothesize that sleep deprivation induced DVB neurite extension is not stabilized, leading to rebalance in extension and retraction, as both processes are affected (retraction by sleep loss). Therefore, DVB morphology is similar to controls and the time to spicule protraction is not increased. Lack of energy conservation due to impaired sleep and potential energy expenditure in the spicule protraction neurons and muscles may lead to further loss of energy, and altered response to sleep deprivation may be detrimental.

The question of *why animals sleep* has yet to be answered; however, there are many supported hypotheses, including energy regulation^105–109^ and neural plasticity^25–28, 38–40, 110–112^. Sleep deprived *C. elegans* undergo less movement quiescence^89, 90, 95^ and likely have increased energy expenditures during a period of energy conservation^92^. Therefore, we propose that structural changes in response to sleep deprivation may prevent circuit processes or behaviors that have higher energetic demands. The spicule protraction circuit becomes progressively hyperexcitable in adult males^103^ and increase in DVB neurite outgrowth likely inhibits the increased muscle excitability and slows spicule protraction, thereby conserving energy. Another proposed role for sleep is in synaptic downscaling, circuit consolidation, and other plasticity mechanisms^27, 45, 110–112^. This role for sleep fits with our hypothesis that DVB morphology is determined by the balance of neurite extension and retraction, where animals with less sleep, and therefore decreased neurite retraction, have overall increased neurite outgrowth and synapses **(Figure 6B)**.

We propose that DVB neurite length in animals lacking *nrx-1* or *nlg-1* at day 1 do not differ from controls, as processes which are not properly stabilized without these essential synaptic adhesion molecules are likely retracted in *C. elegans* during sleep. Our model predicts that in sleep deprived *nrx-1* or *nlg-1* mutant animals, neurites that would normally grow out in response to sleep loss do not do so as they may not be able to be stabilized **(Figure 6C)**. The lack of DVB GABAergic inhibition in *nrx-1* or *nlg-1* after sleep loss also means that they are not able to compensate for energy expenditure during typical sleep periods by slowing spicule protraction and inhibiting circuit excitability **(Figure 6C)**. Therefore, the altered neuronal and behavioral changes after sleep loss in animals without *nrx-1* or *nlg-1* may have detrimental impacts.

Neurexin/*NRXN1* and neuroligin/*NLGN3* are both associated with autism and a number of other neurodevelopmental and neuropsychiatric disorders and conditions^55^. Between 50-85% of individuals with autism report sleep problems compared to 12-40% in neurotypical individuals^21, 23^, underscoring the need for further research into the effect of sleep loss in this condition. Additionally, our identification of sexual maturation as critical for sleep deprivation induced neuroplasticity is particularly interesting given the many connections between neurological conditions and adolescence. In fact, individuals with autism have heightened responses to stress during adolescence and increased variability in cortisol levels, including increases at night^113^. Increased stress reactivity in adolescence may increase the likelihood of comorbid anxiety and sleep disorders^113^. Other neuropsychiatric conditions including schizophrenia (*NRXN1* associated) have an average age of onset in mid to late adolescence or shortly after, suggesting selective vulnerability during this period. Further, the specific involvement of *nrx-1* alpha in regulating structural response to sleep deprivation is particularly relevant as most human NRXN1 mutations affect the long (alpha) isoform^115^.

There is clear evidence for a relationship between sleep and autism; however, the directionality between these two is uncertain. Sleep loss may modify characteristic behaviors of autism (social impairment, repetitive and fixated behavior) and, concurrently, sleep problems may be intensified by alterations in anxiety, sensory sensitivity or integration,^16–24^. Our results identify autism SAM genes that contribute to the response of neuronal circuits and behavior to sleep deprivation through modification of plasticity mechanisms. Therefore, we may expect altered neuronal response to sleep loss in individuals with variants in SAM genes, leading to altered or additional behavioral changes. This is supported by evidence that sleep interventions can modify behavior in autism^23, 24^. Our findings present a model system in which to continue studying the directional relationships between sleep, autism genes, and relevant behaviors.

Here, we provide novel insights into the structural and functional effects of sleep deprivation on neurons that depend on the autism-associated genes neurexin and neuroligin. This work overcomes numerous methodological caveats in the study of sleep dependent plasticity in more complex systems by using four specific and complementary methods to disrupt sleep in *C. elegans.* Additionally, our results integrate analysis of sleep dependent plasticity at every level, including neurons, synapses, and behaviors. These findings highlight the importance of conserved SAMs in the regulation of sleep dependent structural neuroplasticity and the behavioral output of a single neuron. This work provides examination, at an unprecedented resolution, into the interplay of sleep, structural plasticity, neuronal function, and behavior, with potential translational relevance for the impact of sleep disruption in neurodevelopmental conditions.

## ACKNOWLEDGEMENTS

We thank the members of the Hart lab for technical assistance and intellectual input on this project. The authors thank all *C. elegans* labs at the University of Pennsylvania, including the labs of Meera Sundaram, John Murray, Chris Fang-Yen, and Colin Conine for providing feedback on this project. We express appreciation for Anthony D. Fouad from Tau Scientific for technical support on the WormWatcher Platforms. Figures were generated in part with BioRender.com. Some strains were provided by the CGC, funded by NIH Office of Research Infrastructure Programs (P40 OD010440). This work was supported by a SFARI Bridge to Independence Award (MPH), the Autism Spectrum Program of Excellence at the Perelman School of Medicine, the National Science Foundation Graduate Research Fellowship Program (MHC), the 2020 Hearst Foundation Fellowship (MHC), and by NINDS (1R01NS129736(MPH), R01NS107969(DMR), and R01NS122779(DMR)) of the NIH.

## AUTHOR CONTRIBUTIONS

MHC, DMR, and MPH developed the experiments for this study. MHC conducted all imaging and behavioral experiments apart from vibration stimulus performed by MPH. MHC performed all analysis for this project and created all data figures. MHC, DMR, and MPH wrote, reviewed, revised, and approved the manuscript.

## FIGURES

**Supplemental Figure 1.**
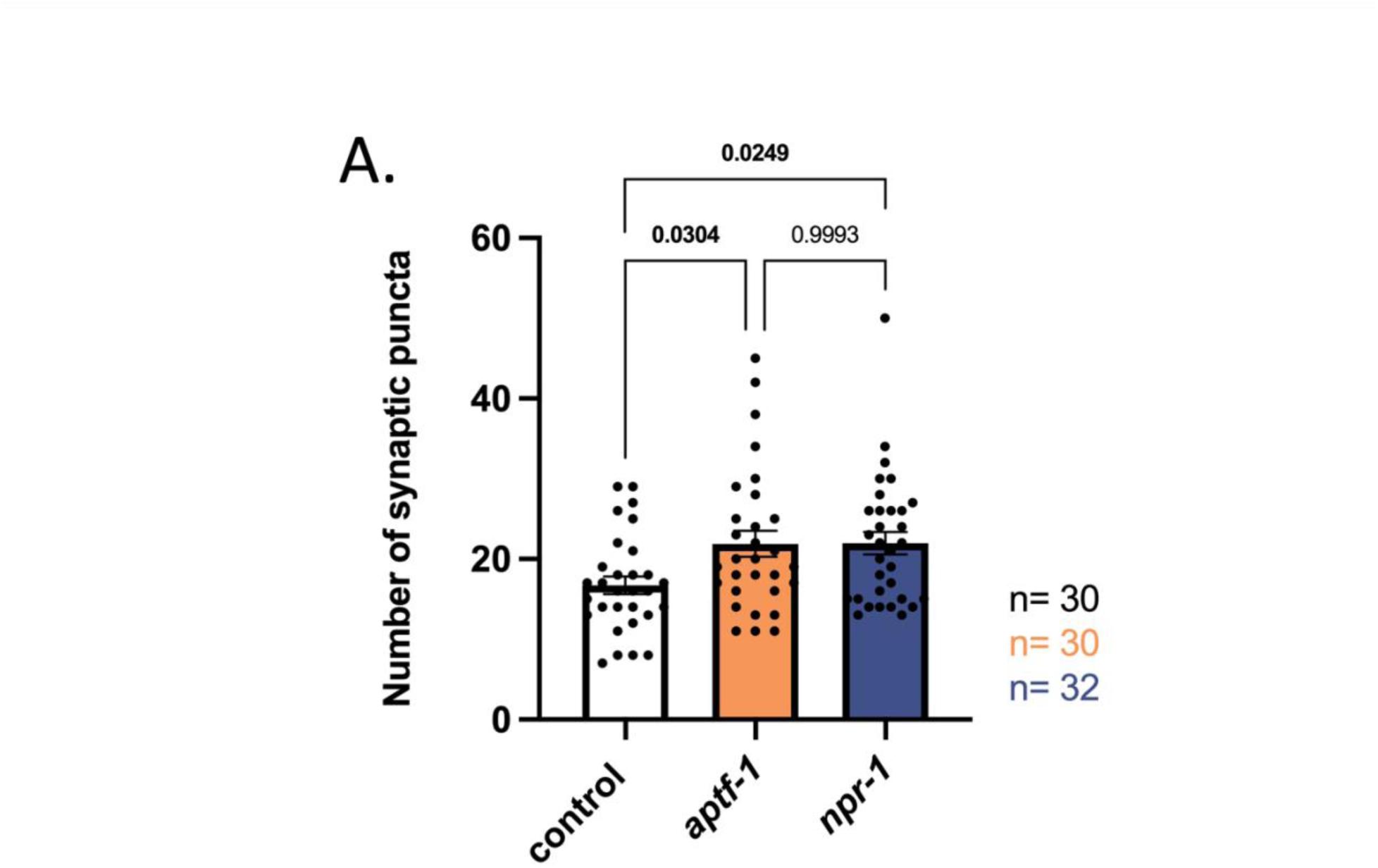
A) Quantification of *cla-1::gfp* puncta area in DVB in controls, sleep deprived *aptf-1*, and sleep deprived *npr-1* males at day 1. Number of individual animals is indicated by “n”, P-values from one-way ANOVA with Tukey’s post-hoc test. ns=not significant.

**Supplemental Figure 2.**
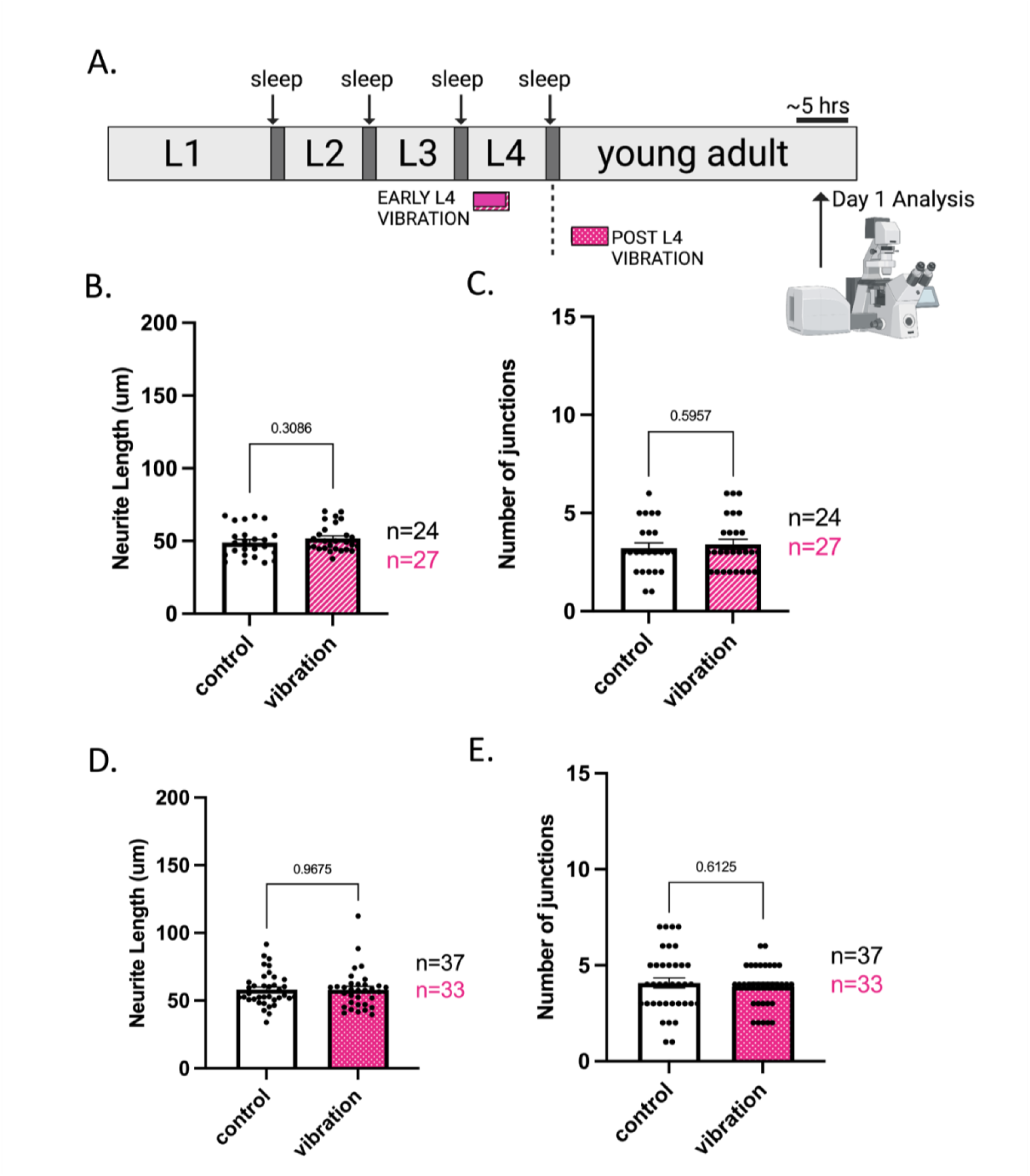
**A)** Schematic of *C. elegans* developmentally timed sleep stages in between each larval stage with timing of vibration application indicated. Striped fill represents early L4 vibration application. Polka dot fill represents post-L4 vibration application. Quantification of DVB **(B)** total neurite length and **(C)** number of junctions in controls and early L4 vibrated males, imaged Number of individual animals is indicated by “n”, P-values from two-tailed unpaired t-test shown.

**Supplemental Figure 3.**
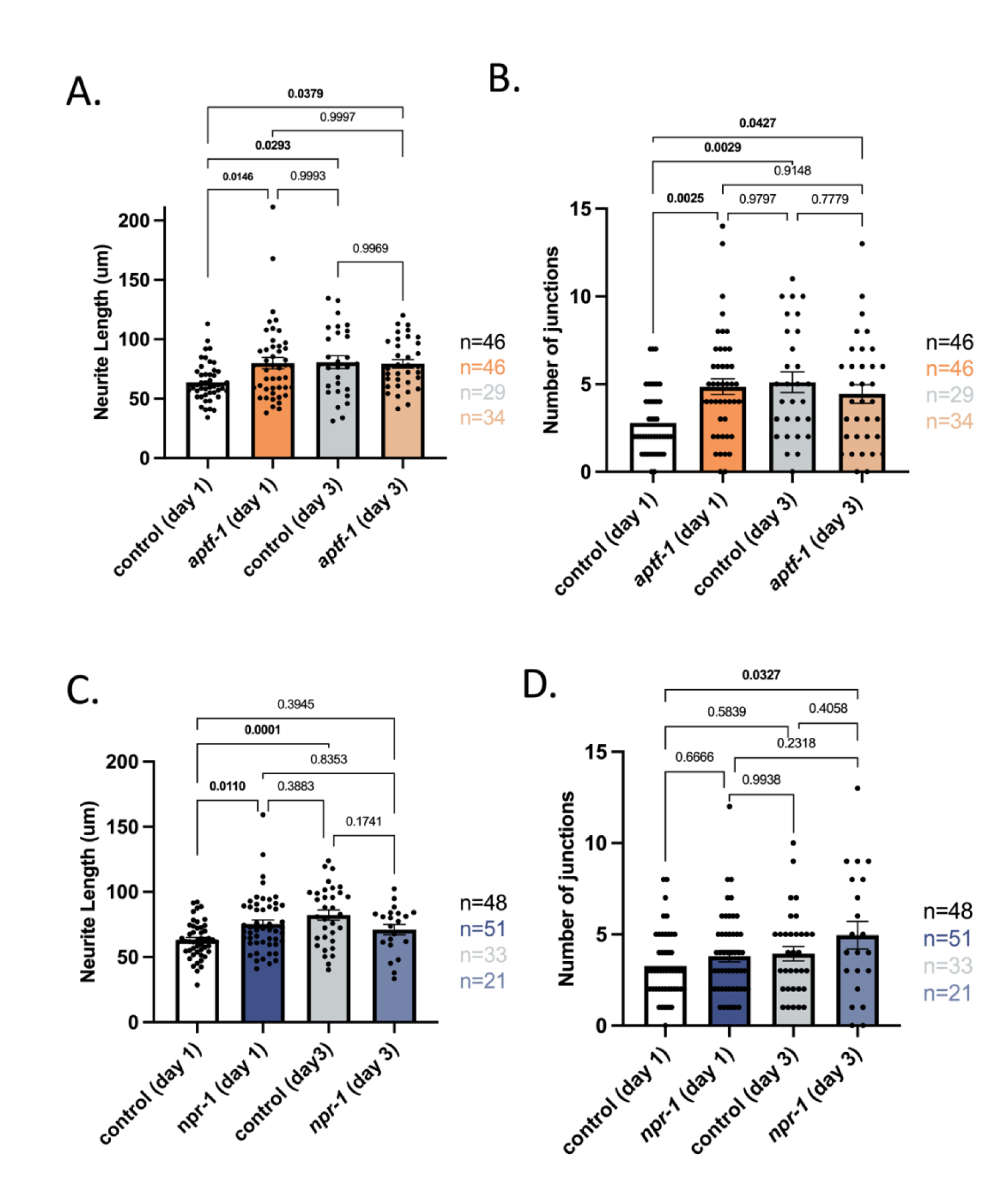
Graph showing **(A)** DVB neurite length and **(B)** junctions in day 1 *aptf-1* males and day 3 controls and day 3 *aptf-1* mutants. **C)** DVB neurite length and **(D)** number of DVB junctions in day 1 *npr-1* males compared to day 3 controls and day 1 *npr-*1 males. Data combined from figure 1 and figure 5. Day 1 and day 3 data collected on separate days. Number of individual animals is indicated by “n”, P-values from one-way ANOVA with Tukey’s post-hoc test shown.

